# Comparing Genomic and Epigenomic Features across Species Using the WashU Comparative Epigenome Browser

**DOI:** 10.1101/2022.11.29.518374

**Authors:** Xiaoyu Zhuo, Silas Hsu, Deepak Purushotham, Samuel Chen, Daofeng Li, Ting Wang

## Abstract

Genome browsers have become an intuitive and critical tool to visualize and analyze genomic features and data. Conventional genome browsers display data/annotations on a single reference genome/assembly; there are also genomic alignment viewer/browsers that help users visualize alignment, mismatch, and rearrangement between syntenic regions. However, there is a growing need for a comparative epigenome browser that can display genomic and epigenomic datasets across different species and enable users to compare them between syntenic regions. Here, we present the WashU Comparative Epigenome Browser (http://comparativegateway.wustl.edu). It allows users to load functional genomic datasets/annotations mapped to different genomes and display them over syntenic regions simultaneously. The browser also displays genetic differences between the genomes from single nucleotide variants (SNVs) to structural variants (SVs) to visualize the association between epigenomic differences and genetic differences. Instead of anchoring all datasets to the reference genome coordinates, it creates independent coordinates of different genome assemblies to faithfully present features and data mapped to different genomes. It uses a simple, intuitive genome-align track to illustrate the syntenic relationship between different species. It extends the widely used WashU Epigenome Browser infrastructure and can be expanded to support multiple species. This new browser function will greatly facilitate comparative genomic/epigenomic research, as well as support the recent growing needs to directly compare and benchmark the T2T CHM13 assembly and other human genome assemblies.

## Introduction

To meet the need to visualize genomic sequences and features at different scales in the genomic era, scientists developed genome browser/viewers to help interpret genomes. The UCSC Genome Browser, equipped with comprehensive annotations and intuitive navigation, gained widespread popularity in the community (Kent et al. 2002; Lee et al. 2022). In addition to the UCSC Genome Browser, there are multiple other tools available to visualize genomes each with its own advantages and focuses (e.g., Ensembl (Fernández-Suárez and Schuster 2010; Cunningham et al. 2022), GBrowse (Stein et al. 2002), WashU Epigenome Browser (Li et al. 2019, 2022; Zhou et al. 2011), IGV (Robinson et al. 2011, 2022), and JBrowse (Buels et al. 2016; Diesh et al. 2022)).

With sharply decreasing sequencing cost, many more genomes of different species become available, and there is an increased effort around the world to systematically sequence a wide variety of organisms (Rhie et al. 2021; Feng et al. 2020; Teeling et al. 2018). The advancement in sequencing technology also promoted many functional genomic assays, which enabled functional annotation of genomic regions (ENCODE Project Consortium 2012; Roadmap Epigenomics Consortium et al. 2015; Dekker et al. 2017; Bujold et al. 2016). Based on whole genome alignment between species, orthologous regions can be directly compared, and insights on conservation and adaptation of genomic features can be drawn. Comparative genomics thus has become an important tool to decipher genomic code (Alföldi and Lindblad-Toh 2013). Comparative epigenomics, which compares the epigenomic features of orthologous regions of multiple species, is also gaining popularity (Xiao et al. 2012; Prescott et al. 2015; Zhou et al. 2017; Modzelewski et al. 2021).

Starting from Miropeats, various visualization tools have been developed to display regional orthologous relationship between species (Parsons 1995; Guy et al. 2010; Sullivan et al. 2011; Goel and Schneeberger 2022; Vollger 2022; dporubsky 2021). These tools provide a variety of comparative features. The gEVAL Browser was designed for genome assembly quality evaluation and can be used to visualize and compare genome assemblies (Chow et al. 2016). Nguyen et al. developed comparative assembly hubs using UCSC Genome Browser’s framework (Nguyen et al. 2014). It utilizes snake track to show multiple query assemblies aligned to a target assembly, and annotations mapped to query assemblies can also be displayed with an automatic “liftOver”. JBrowse2 v1.6.4 also starts to support cross-species comparison in synteny views (Buels et al. 2016; Diesh et al. 2022). CEpBrowser was developed to compare epigenomic datasets between human, mouse, and pig based on the UCSC Genome Browser framework in a gene-centric manner (Cao and Zhong 2013). It organizes linear representation of different species in different windows parallelly. By displaying different species in different windows, CEpBrowser can be implemented relatively easily without breaking the continuity of each genome. However, it only marks syntenic regions using the same color scheme but does not connect syntenic regions from different species or display any genetic differences. In addition, only comparisons between human (hg19), mouse (mm9) and pig (susScr2) are supported. Despite being the first comparative epigenome browser, it has not been widely used by the scientific community.

The WashU Epigenome Browser was developed in 2010 to host and display massive epigenomics datasets (Zhou et al. 2011; Li et al. 2019, 2022). It hosts datasets generated from Roadmap Epigenomics Project (Roadmap Epigenomics Consortium et al. 2015), ENCyclopedia Of DNA Elements (ENCODE) (ENCODE Project Consortium 2012), International Human Epigenome Consortium (IHEC) (Bujold et al. 2016), The Cancer Genome Atlas (TCGA) (Hutter and Zenklusen 2018), Toxicant Exposures and Responses by Genomic and Epigenomic Regulators of Transcription (TaRGET) (Wang et al. 2018), and 4D Nucleome Project (4DN) (Dekker et al. 2017). We recently refactored the browser and vastly improved its performance (Li et al. 2019, 2022).

Build upon the WashU Epigenome Browser, we developed the WashU Comparative Epigenome Browser based on four principles: 1, each assembly uses its own coordinates to anchor annotation and datasets mapped to it; 2, orthologous relationship and genetic variations between assemblies are intuitively illustrated; 3, adaptable to display any whole genome alignment at different scales and resolution; 4, inherits all features of modern genome browsers to facilitate user experience. Here we present the WashU Comparative Epigenome Browser to address the needs to navigate multiple genomes at once and visualize comparative genomics/epigenomics data.

## Results

### The genome-align track connects syntenic regions of two genome assemblies

The foundation that enables comparative genome browsing is the alignment between genome assemblies. We developed a new track type called “genome-align track” which contains genome-wide syntenic relationship between the reference (target) genome and the secondary (query) genome at base-pair resolution. The genome-align track file can be constructed from standard chained alignment AXT files (Schwartz et al. 2003) using customized tools we developed.

We created a comparative epigenome gateway to help organize and facilitate the selection and display of curated genome-align tracks (http://comparativegateway.wustl.edu/). Using this gateway, the users first select the reference assembly. When one reference genome is selected, all the available genome-align tracks will be populated as a list of secondary genomes (Fig. 1). Then the user can select one or more genome-align tracks anchored to the reference genome, save the selection, and open a new WashU Epigenome Browser window with all the selected genome-align tracks. With genome-align tracks loaded, the user can then use the browser’s web interface to load available annotations (Tracks -> Annotation Tracks), public data (Tracks -> Public Data Hubs), or user’s own data (Tracks -> Remote/Local Tracks) on the browser mapped to either reference genome or any of the loaded secondary genomes (Fig. 1).

**Figure 1:**
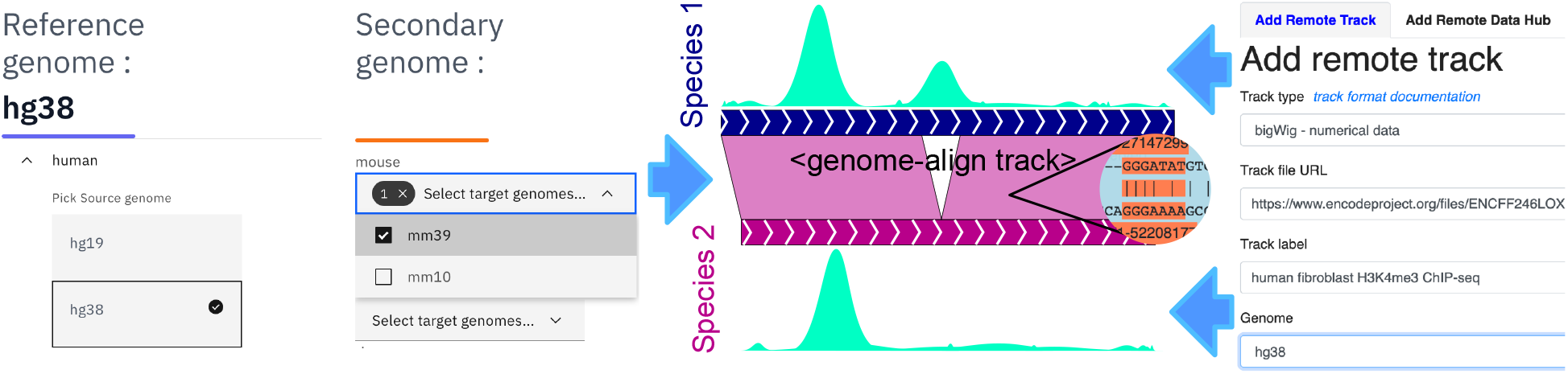
The web user interface of the WashU Comparative Epigenome Browser. Genome-align track selector web interface is shown on the left. After selecting desired alignment tracks, the user will be redirected to the main WashU Epigenome Browser. At last, the user can load data and annotations to either reference or secondary genomes on the main browser site.

The genome-align track supports comprehensive, multi-resolution genome alignment display. At the finest resolution, orthologous coordinates from query genomes are vertically aligned and anchored to the reference genome. Detailed whole-genome alignment at the single nucleotide resolution is displayed in the genome-align track, enabling users to navigate and examine the genetic differences between the query genome and the reference genome. It is straightforward to visualize single-nucleotide variations (SNVs) and short insertion/deletions (indels) between the two genome assemblies (Fig. 2a).

**Figure 2:**
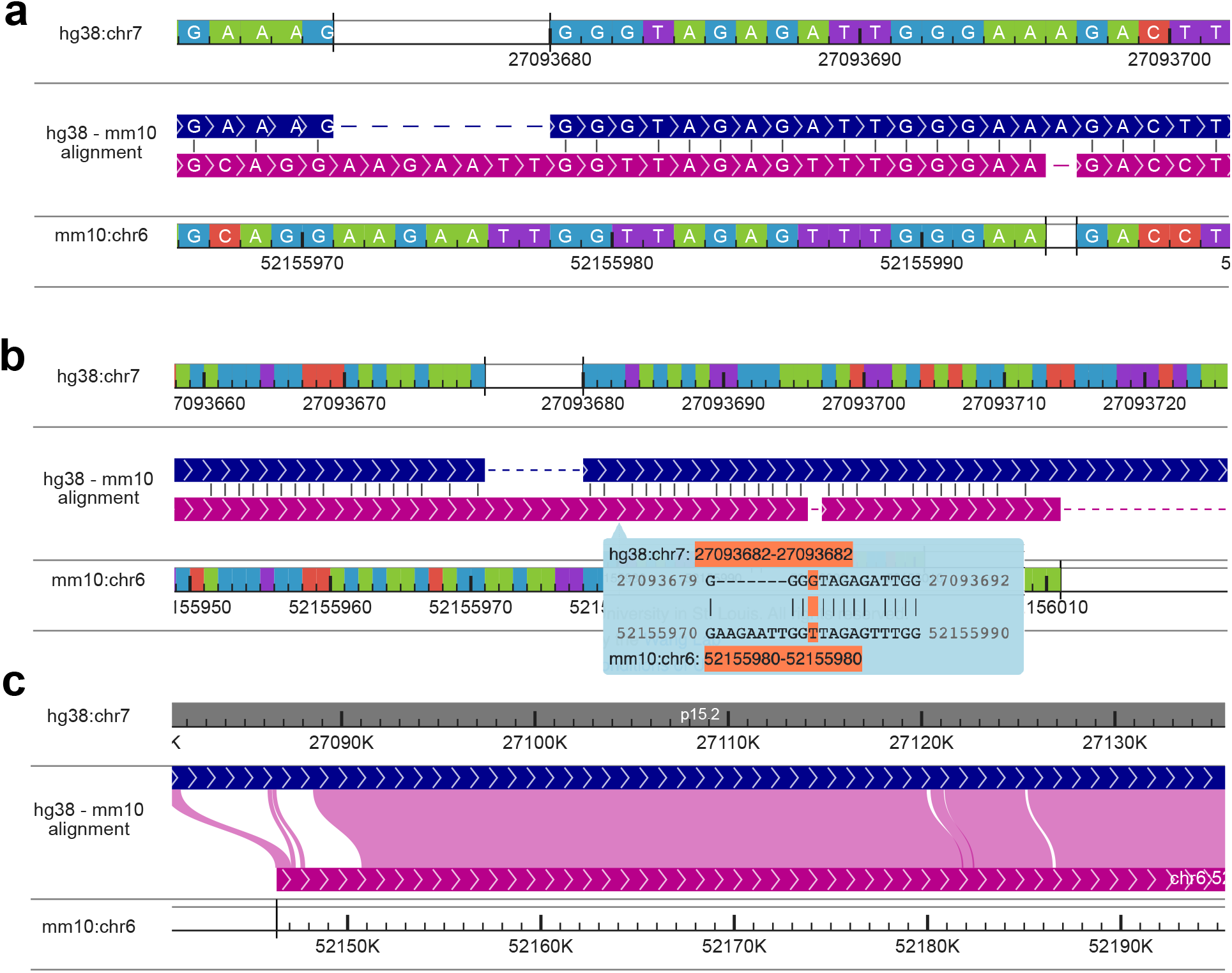
Display genome alignment using the WashU Comparative Epigenome Browser. a: Displaying hg38-mm10 blastz alignment at the nucleotide level with > 10 pixels per nucleotide. Sequence strand in the alignment is illustrated using arrows. Syntenic nucleotides from hg38 and mm10 are vertically aligned with gaps inserted. Same nucleotides are illustrated by a short vertical line. b: Displaying hg38-mm10 alignment between 0.1 pixels per nucleotide and 10 pixels per nucleotide. The alignment is organized the same as panel A without displaying nucleotides within the alignment. Alignments at nucleotide resolution are visible in the cursor tip hover box and the nucleotide alignment under the cursor is highlighted in orange (G - T). c: Displaying alignment with > 10 nucleotides per pixel. Both hg38 and mm10 genomes are continuously displayed without gaps. Syntenic regions are connected using Bezier curves.

Users can pan and zoom on the genome-align track using the tools bar on top of the displayed window in a similar fashion as they operate on any other browser track types. When the number of nucleotides within a browser window exceeds the available pixels to display each nucleotide clearly (10 pixels per nucleotide), the browser stops displaying individual nucleotides within the alignment. Instead, it would display a 20-bp alignment in a floating box next to the cursor when the user mouses over the genome-align track (Fig. 2b). This feature helps users to visualize a larger aligned region without missing the base-pair resolution information in the alignment. Vertically aligning and anchoring query genomes to the reference genome is a straightforward and convenient way to display SNVs and small indels between query and reference genomes. However, it is insufficient to show any large, more complexed structural variations (SVs) between species. The WashU Comparative Epigenome Browser displays both the reference and query genomes in a linear manner and connect syntenic regions using Bezier curves if the browser window contains a long genomic alignment (more than 10 bases per pixel) (Fig. 2c). By doing so, large scale genetic variations can be directly visualized in the browser. Since both genomes are continuously and co-linearly displayed, epigenomic features are also displayed in full without sudden truncation.

### Using the WashU Comparative Epigenome Browser to compare epigenomic features between species

The genome-align track is more than just a visualization tool to display pairwise whole-genome alignments. After loading the genome-align track onto the browser, users can load annotations and datasets mapped to the secondary genome in the browser and compare them with those mapped to the reference genome. With this feature, the browser connects annotations and datasets from different genomes together using their syntenic relationship in the same window. While users navigate the reference genome, the browser retrieves syntenic coordinates from other genomes and fetches all the loaded tracks.

We can use the browser to characterize deeply conserved epigenomic marks. In Fig. 3a the browser displays deeply conserved CpG methylation in liver between mouse and zebrafish using methylC tracks (Yue et al. 2014; Yang et al. 2020; Zhou et al. 2014). By displaying the Hox C gene cluster from both mouse and zebrafish reference genomes and their syntenic relationship, we can appreciate that only a small fraction of their genomic sequences can be aligned with each other after hundreds of million years of independent evolution, recapitulating the discovery made by Zhang et al. (Zhang et al. 2016). Even conserved CpG islands between these two species are sparse. However, except for a few species-specific transposable elements, the CpG sites are hypomethylated in the region in both species. Despite limited sequence conservation, the apparent epigenomic conservation suggests deeply conserved regulatory pattern in the region.

**Figure 3:**
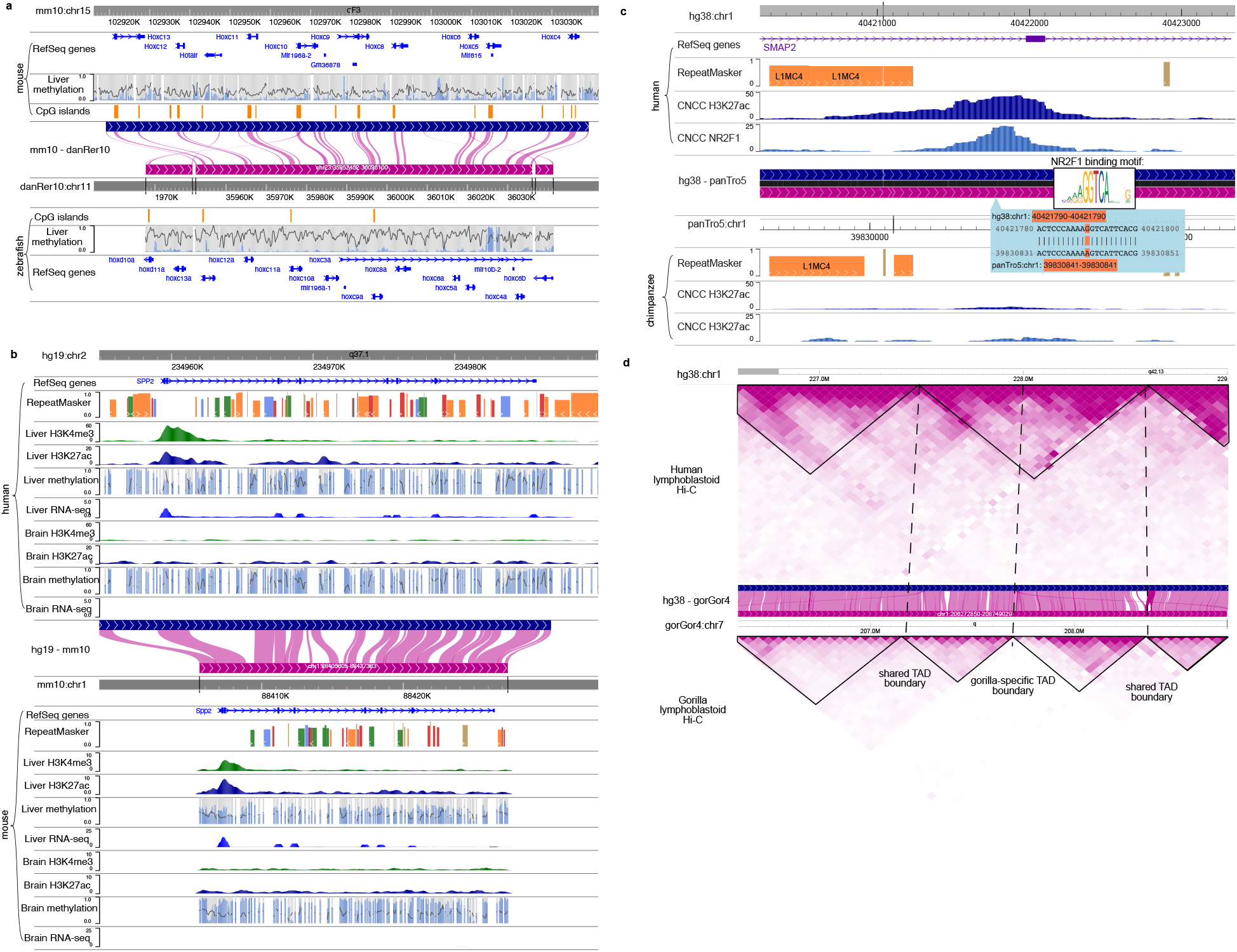
Compare epigenomes between species. a: The DNA methylation status of Hox C gene cluster is conserved between mouse and zebrafish. Mouse and zebrafish DNA methylomes were characterized by Zhang et al. Mouse and zebrafish reference genome (mm10 and danRer7) are shown back-to-back anchored by the mouse-zebrafish genome-align track along with their gene, repeat, and CpG Island annotations. Liver DNA methylome data from Zhang et al. using Enhanced Reduced Representation Bisulfite Sequencing (ERRBS) (Zhang et al. 2016) is displayed. b: H3K4me3 and H3K27ac ChIP-seq, WGBS and RNA-seq of brain and liver samples from both human and mouse of SPP2/Spp2 gene are displayed using the WashU Comparative Epigenome Browser. Both DNA methylation level and read depth are illustrated in the MethylC track. c: Lineage-specific epigenomic innovation. H3K27ac, NR2F1 ChIP-seq data from both human and chimpanzee CNCC in SMAP2 gene regions were plotted in the WashU Comparative Epigenome Browser. A human-specific NR2F1 and H3K27ac peak suggests a putative human-specific enhancer in this region. The putative enhancer is associated with a human-specific NR2F1 binding motif. d: 3D genome structure differences between species. Human lymphoblastoid Hi-C contact map mapped to hg38 and gorilla lymphoblastoid Hi-C data mapped to gorGor4 were compared by anchoring to the human-gorilla alignment track.

Epigenomic modifications underlie tissue specificity. It has been shown before that the tissue-specific epigenomic patterns are often conserved between species (Zhou et al. 2017). The comparative browser makes it intuitive to examine the conservation pattern of tissue-specific gene activities. Fig. 3b illustrates the conserved liver-specific expression and epigenome landscape of gene Secreted Phosphoprotein 2 (SPP2) between human and mouse. Epigenomic data, including Whole-Genome Bisulfite Sequencing (WGBS), H3K4me3 ChIP-seq, H3K27ac ChIP-seq, and RNA-seq data of liver and brain from ENCODE and mouse ENCODE are displayed on the respective reference genomes in the comparative browser (ENCODE Project Consortium 2012; Yue et al. 2014) spanning the syntenic region around human SPP2 gene and its orthologous mouse Spp2 gene (Fig. 3b). Both species share the pattern of liver-specific active histone marks, low DNA methylation and high RNA expression, as well as lack of active histone/expression and high DNA methylation in the brain, indicating epigenetic conservation.

In addition to showcasing conserved features, the browser is equally effective at visualizing lineage-specific epigenomic features. Fig. 3c displays H3K27ac and transcription factor NR2F1 ChIP-seq data from iPSC-derived Cranial Neural Crest Cells (CNCC) of both human and chimpanzee (Prescott et al. 2015). This region contains a putative human-specific enhancer, defined by the co-occurrence of NR2F1-binding and H3K27ac peak in the intron of SMAP2 gene. The epigenomic signature suggests that this is either a human-gain or chimpanzee-loss of a putative CNCC enhancer. Zooming in to examine the alignment at base level, we identified a single nucleotide difference between human and chimpanzee that maps to a high information content position in the NR2F motif, potentially explaining the enhancer gain or loss. This example demonstrates that our browser can be used to associate epigenomic differences between species with their genetic differences.

The comparative browser also supports visualization and comparison of long-range chromatin interaction data across different genomes, thus facilitating the studies of 3D genome evolution (Vietri Rudan et al. 2015). Fig. 3d directly compares the 3D genome structure between human and gorilla in the human chr1 q42.13 region. Hi-C data from lymphoblastoid cells of human and gorilla reveals several conserved TADs. Interestingly, one TAD in human is split into two different TADs in the gorilla. This observation using the comparative browser recapitulated insights from Yang et al. (Yang et al. 2019).

### Visualizing the relationship between genomic variation and epigenomic variation

There has been a growing interest in understanding the relationship between genetic variation and epigenetic variation. We have already demonstrated using the browser to display the association between epigenomic changes with a SNP (Fig. 3c). Recently we characterized structural variations (SVs) between human and chimpanzee and their impact on the epigenome (Zhuo et al. 2020). Fig. 4a illustrates an interesting case of human-specific TE-derived putative enhancer. In this comparative browser view, investigators can easily and intuitively compare a species-specific TE insertion and its associated epigenomic modification. Here, a human-specific retrotransposon SVA_F appears in the intron of the DNMBP gene. The sequence of this SVA_F element is highly repetitive, thus it exhibits low mappability scores (average 50bp score <0.05) indicating that short sequencing reads derived from this element may not be uniquely mapped back (Derrien et al. 2012). Indeed, a cranial neural crest cell (CNCC) H3K27ac ChIP-seq dataset (sequenced using 50 bp reads) does not contain signal within the SVA_F element but reveals a peak at the 3’ boundary of the element. Further analysis suggests that this boundary peak reflects enhancer signals from within this SVA_F element (Zhuo et al. 2020). In contrast, an iPSC H3K9me3 ChIP-seq dataset (sequenced using 100 bp paired-end reads) is able to uniquely reveal an enrichment peak over this SVA_F element, indicating the deployment of repressive chromatin onto this newly inserted retrotransposon in iPSC (Zhuo et al. 2020). The parallelly displayed chimpanzee genome and corresponding epigenomic datasets illustrate the lack of this specific SVA_F insertion and absent of respective epigenomic marks. This direct visual comparison of the retrotransposon insertion and epigenomic changes between the two species recapitulates the discovery of a tissue-specific enhancer derived from a human-specific retrotransposon insertion.

**Figure 4:**
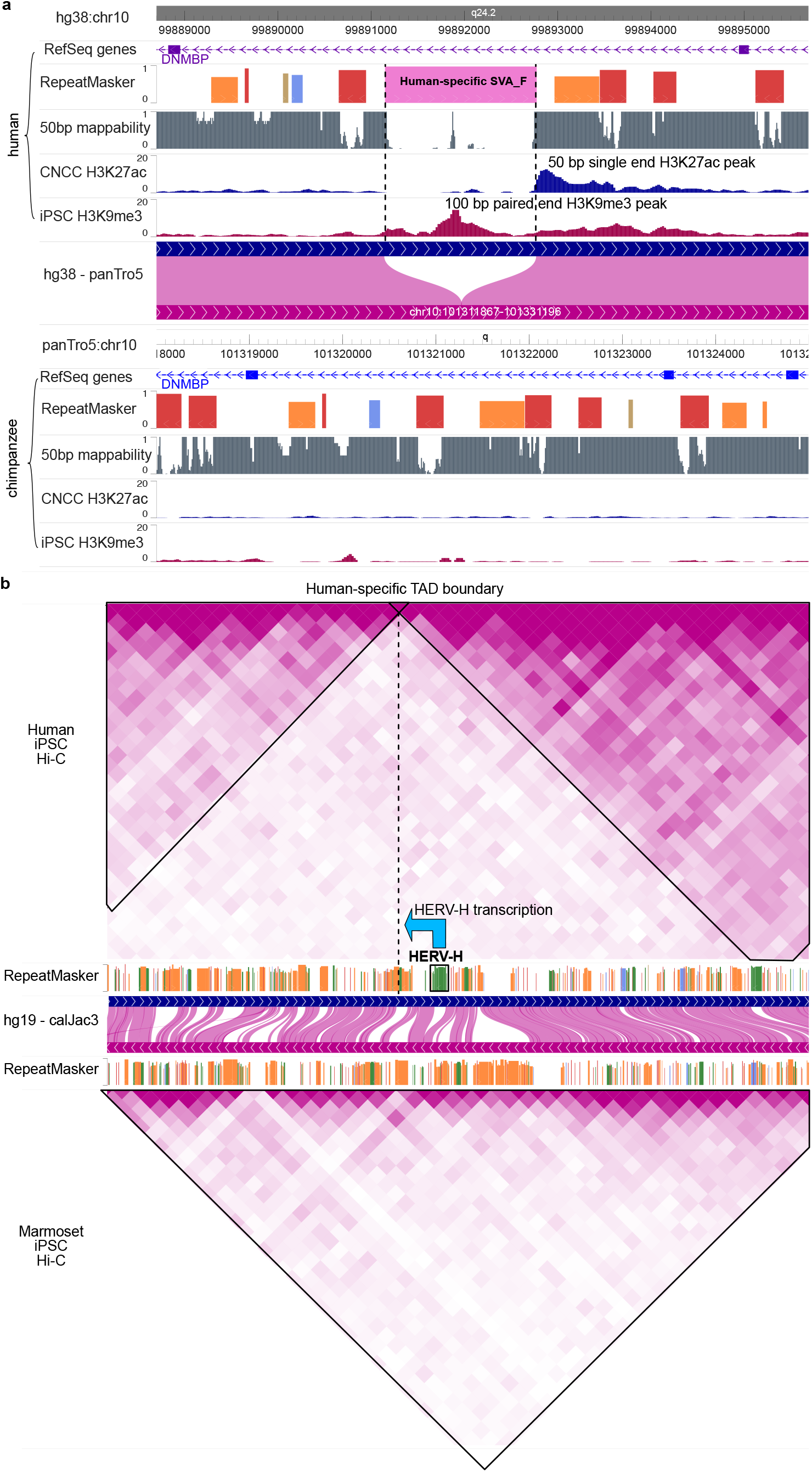
Connecting epigenomic changes with genomic changes using the WashU Comparative Epigenome Browser. a: RefSeq Genes, repeatMasker and 50bp mappability annotations along with H3K27ac ChIP-seq data from Cranial neural crest cell (CNCC) and H3K9me3 ChIP-seq data from iPSC in both human and chimpanzee were plotted in DNMBP gene region. The H3K9me3 peak in the human-specific SVA insertion indicates epigenomic repression of this element in iPSC and the human-specific H3K27ac and NR2F1 peak indicates the creation of a putative new CNCC enhancer in the human-lineage. b: Human-specific HERV-H expression is correlated with a new TAD boundary in iPSC in the human genome compared with the marmoset genome.

Zhang et al. demonstrated that the expression of HERV-H is associated with new TAD boundaries in primates (Zhang et al. 2019). This association can be easily appreciated in a comparative browser view. In Fig. 4b Hi-C maps of human iPSC and marmoset iPSC can be directly compared in the context of their genome alignment. In the human genome, an HERV-H insertion is associated with a human-specific TAD boundary reflected by the Hi-C contact map, which is absent in the marmoset genome (Fig. 4b). It is notable that the TAD boundary is ∼20kb downstream of the HERV-H insertion in the human genome, suggesting it is the expression instead of the presence of binding motif in the HERV-H that contributes to the TAD boundary. These examples demonstrate that the WashU Comparative Epigenome Browser can be used to directly compare genomic datasets across species and visualize the association with genetic changes.

### Displaying genome annotations and datasets from multiple species using the WashU Comparative Epigenome Browser

A natural extension of the pairwise comparison functions is to support comparison among multiple species. Conceptually, this extension is equivalent to visualizing genomic data aligned to a multiple genome alignment across species. Practically, we use multiple genome-align tracks to anchor the visualization to the same reference genome, thus enabling an intuitive comparison of genomic data across orthologous regions of multiple species. We use CTCF turnover events characterized in Schmidt et. al. and Choudhary et. al. (Schmidt et al. 2012; Choudhary et al. 2020) to illustrate the comparative analysis across multiple genomes.

Schmidt et. al. characterized the CTCF binding sites of six mammalian species (human, macaque, mouse, rat, dog and opossum) and identified thousands of conserved as well as lineage-specific, retrotransposon-derived CTCF binding sites (Schmidt et al. 2012). We display both CTCF ChIP-seq data and called CTCF binding peaks of the six species from this study using the WashU Comparative Epigenome Browser, anchored on the human reference genome hg19. (Fig. 5). This allows direct comparison of CTCF binding across species along with genetic changes in each lineage.

**Figure 5:**
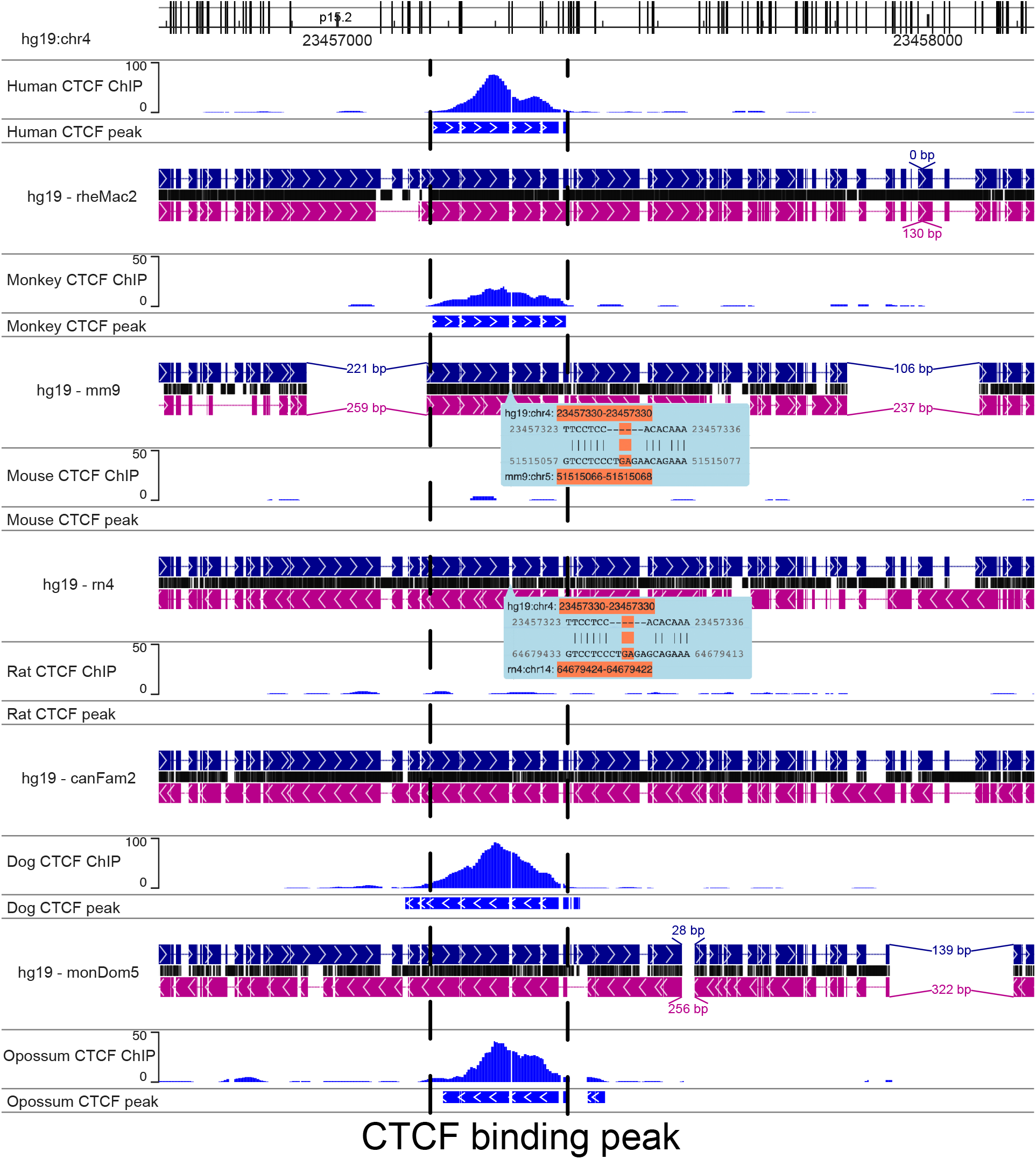
Using the WashU Comparative Epigenome Browser to visualize and compare the CTCF binding sites from six mammals. Genes, repeats, CTCF ChIP-seq and input from human (hg19), rhesus macaque (rheMac2), mouse (mm9), rat (rn4), dog (canFam2) and opossum (monDom5) were displayed on the browser. Human reference genome hg19 was used as the reference genome and all the other species were anchored to their orthologous region from hg19 using whole genome alignments. A: hg19: chr4:23456625-23458090 region shows a conserved CTCF binding peak in the orthologous loci in all mammal genomes except the two rodents, indicates a rodent-specific loss of a conserved CTCF binding site. The loss of CTCF binding is also coincided with a rodent-specific 6bp insertion.

Fig. 5 highlights the loss of a conserved CTCF binding sites in rodents (Fig. 5). All genome assemblies are vertically aligned, and interruptions are introduced in the tracks when gaps occur in either reference or secondary genomes. In contrast to the other four genomes, mouse and rat do not display a CTCF binding peak in this region, and this event is associated with a rodent-specific 6 bp insertion in the ortholog site of the CTCF site conserved in the other four species. Again, the WashU Comparative Epigenome Browser makes it intuitive to display and identify associations between genetic changes and epigenomic changes across multiple species.

### Extending comparative genomic analysis to non-model organisms and new assemblies

The WashU Comparative Epigenome Browser is built on an actively maintained and expandable platform. New genomes are routinely added to the browser to serve scientists around the world. The browser engineers respond to new comments and feature request (including request for new genomes) on the browser GitHub repository frequently (https://github.com/lidaof/eg-react/issues). We also documented how to add new genomes to the browser for a local environment for advanced users with JavaScript background (https://epigenomegateway.readthedocs.io/en/latest/add.html).

Using this flexible framework, we created multiple non-model organism reference genomes in our browser. For example, we created reference cattle genome UMD_3.1.1/bosTau8, and generated bosTau8-mm10 genome-align track using bosTau8 as the reference genome. Fig. 6a displays a direct comparison of DNA methylation patterns between cattle and mouse across heart, lung and liver (Liu et al. 2020; Zhou et al. 2020). We display the methylation pattern of liver-specific gene Spp2 promoter in the comparative browser, and we can see the tissue-specific methylation pattern is conserved between mouse and cow (Fig. 6a). Thus, the application of the WashU Comparative Epigenome Browser can easily extend beyond traditional model organisms.

**Figure 6:**
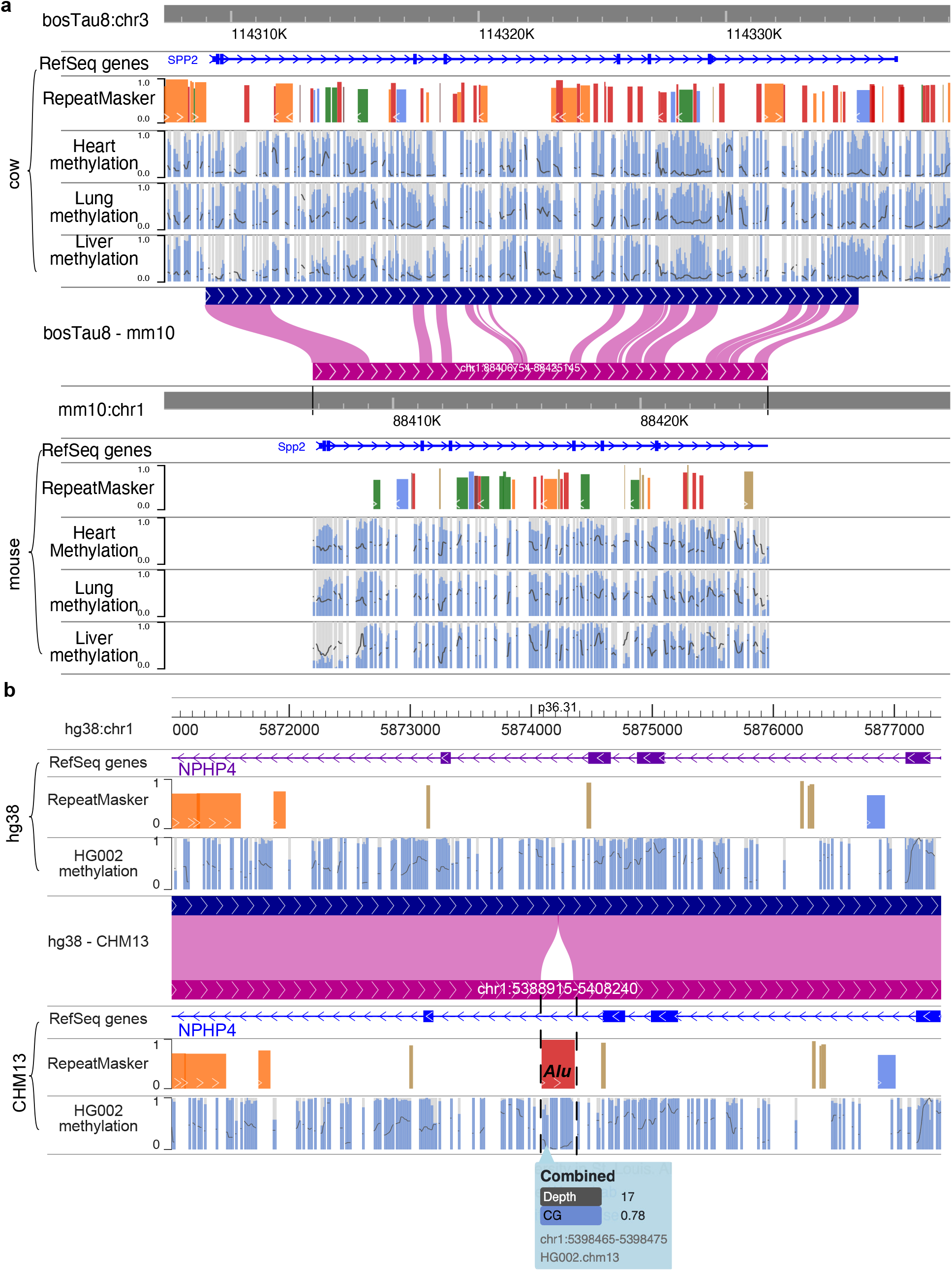
a: Create a cattle-mouse comparative browser view and use it to compare DNA methylation in heart, lung and liver between cow and mouse. RefSeq genes, repeatMasker tracks along with DNA methylation status of heart, lung and liver tissues from both cow and mouse were plotted on the Comparative Epigenome Browser. b: Utilizing the browser to compare the difference between hg38 and CHM13 and how it may affect genomic analysis. The same HG002 WGBS data was mapped to hg38 and CHM13, respectively. The DNA methylation difference by either genome is minimum across most of the genomic region, but an *Alu* insertion is only presents in the CHM13 reference, and the hypermethylation of this Alu element can only be assessed using the CHM13 reference.

Finally, the comparative browser also fulfills a growing need in the field to compare and benchmark the performance of different human genome assemblies (Aganezov et al. 2022). The recent release of the T2T CHM13 genome assembly as well as multiple alternative human genome assemblies from the Human Pangenome Reference Consortium (Cheng et al. 2021; Ebert et al. 2021; Jarvis et al. 2022; Wang et al. 2022; Garg et al. 2021; Porubsky et al. 2021) represents a major improvement for genomics, but the impact of analyzing functional genomics data using different genome assemblies remains to be evaluated. Our Browser support direct visualization of such evaluations. We mapped the public HG002 WGBS data (Baid et al. 2020) to both hg38 and CHM13 reference genomes, and in Fig. 6b we illustrate an Alu insertion present in CHM13 but absent in hg38. In this case, the presence and hypermethylation of the Alu in HG002 is only visible when the reads were mapped to the CHM13 reference genome (Foox et al. 2021; Nurk et al. 2021). Therefore, the WashU Comparative Epigenome Browser provides a near-term, conventional visualization of differential mapping results before the maturation of pangenome graph mapping and subsequent visualization (Miga and Wang 2021; Wang et al. 2022; Liao et al. 2022; Hickey et al. 2022; Guarracino et al. 2021).

## Discussion

Here we present the WashU Comparative Epigenome Browser to visualize comparative genomic/epigenomic features. The browser functions may help scientists interested in comparative genomics/epigenomics to examine their regions of interest and produce publication quality browser views to showcase their findings. In addition to a growing number of genomes, genome-align tracks, and genomics datasets we currently host, users can build and host their own comparative browser with customized species and genome builds. It enables scientists, especially those working on non-model organisms, to visualize and compare genomic and epigenomic features of different species.

The comparative features are fundamentally enabled by the genome-align track, a pairwise genomic alignment track derived from AXT format (Schwartz et al. 2003). Comparison across multiple genomes is achieved by using multiple genome-align tracks anchoring to the same reference genome. While it is possible to generalize the comparative functions based on a multi-genome alignment, the pairwise comparison is more technically practical and intuitive on a two-dimension computer screen. We envision continued exploration of advanced web technologies to further enhance the performance of multi-genome comparison.

## Acknowledgement

We want to thank Jian Ma, Yang Yang from Carnegie Mellon University for providing the human and chimpanzee Hi-C dataset.

